# Developing an Antiviral Peptides Predictor with Generative Adversarial Network Data Augmentation

**DOI:** 10.1101/2021.11.29.470292

**Authors:** Tzu-Tang Lin, Yi-Yun Sun, Wei-Chih Cheng, I-Hsuan Lu, Shu-Hwa Chen, Chung-Yen Lin

**Author notes:** Joint first author (Tzu-Tang Lin, Yi-Yun Sun).

## Abstract

**Motivation:** New antiviral drugs are urgently needed because of emerging viral pathogens’ increasing severity and drug resistance. Antiviral peptides (AVPs) have multiple antiviral properties and are appealing candidates for antiviral drug development. We developed a sequence-based binary classifier to identify whether an unknown short peptide has AVP activity. We collected AVP sequence data from six existing databases. We used a generative adversarial network to augment the number of AVPs in the positive training dataset and allow our deep convolutional neural network model to train on more data.

**Results:** Our classifier achieved outstanding performance on the testing dataset compared with other state-of-the-art classifiers. We deployed our trained classifier on a user-friendly web server.

**Availability and implementation:** AI4AVP is freely accessible at http://axp.iis.sinica.edu.tw/AI4AVP/

**Supplementary information:** Supplementary data is also available.

## Introduction

Recently, viral pathogens have become more prevalent and diverse; they pose increasingly severe threats to people. Treating viral diseases clinically with existing antiviral drugs and interferons is inefficient because of side effects and increasing cases of drug resistance (1). Antiviral peptides (AVPs) are characterized by their high specificity and effectiveness against reemerging and drug-resistant viruses, such as coronaviruses (2), through blocking the virus from entering the host cell or inhibiting virus replication. Additionally, natural AVPs have low toxicity and peptidase biodegradability compared with available antiviral drugs (3). Therefore, AVPs are potential candidates for antiviral drug development.

Although several AVP predictors already exist (**Supplementary Table 1**), they were trained using datasets from AVPpred collected in 2012. Recently, many AVPs have been discovered or synthesized, and added to databases. We collected data from these databases and used them to train a prediction model. To increase the size and balance between the positive and negative datasets, we utilized a generative adversarial network (GAN) in the data augmentation process, generating positive data based on real AVPs. We detailed the process of using GAN to generate peptides in a previous study (4) (**Supplementary Figure 1**).

We used PC6 encoding(5), a protein-encoding method based on six physicochemical properties, to transform sequential data into matrices (**Supplementary Figure 2**). Finally, we constructed our prediction model using a convolutional neural network (CNN; **Supplementary Figure 3**).

## Materials and methods

### Data collection and preprocessing

We collected positive AVP data from six existing AVP databases: APD3, DRAMP, YADAMP, DBAASP, CAMP, and AVPdb (**Supplementary Table 2**). We excluded duplicate data and peptides with unusual amino acids (“B,” “Z,” “U,” “J,” “O,” “X,” “i,” “n,” and “−”) and selected AVPs with a length of 10 to 50. We obtained data on 2,934 AVPs from six databases. We used 90% of the data (2,641 AVPs) for training and 10% of the data (293 AVPs) for testing. To ensure that the evaluation of the model’s performance compared with other predictors was fair, we excluded data from the training datasets of those predictors (2012 dataset) in our testing dataset. We combined peptides collected from the Swiss-Prot database and randomly generated sequences to form the negative set. Specifically, we obtained 8,592 peptides from Swiss-Prot with a length between 10 and 50 without AMP-related keywords, such as “antimicrobial,” “antibiotic,” “amphibian defense peptide,” or “antiviral protein.” Subsequently, we randomly arranged 20 amino acids to generate 8,592 sequences to construct a negative dataset with 17,184 data points. Similar to constructing the positive dataset, we randomly selected 293 data points for testing to balance the positive and negative testing datasets.

### Data augmentation by GAN

Balancing the amount of data is essential for training the classification model to avoid biases caused by imbalance between positive and negative labels. In our case, the most direct method of balancing the datasets was to select only 2,641 data points from the negative dataset to fit the positive dataset. However, this would have made the remaining negative data useless for model training. To fully utilize all the negative data, we augmented the amount of positive data using a GAN. The GAN was trained using all AVP data as the input; it then generated much AVP-like data. We added the generated AVP-like data to the original AVP data to achieve parity between the positive and negative datasets. Therefore, we eventually obtained 16,995 positive and 16,995 negative data points for classifier training.

### Protein-encoding method

We used PC6 encoding, a protein-encoding method developed in our previous study, to transform peptide sequence data into matrices. PC6 encoding can consider peptides’ order and the physicochemical properties of amino acids and extract essential features for model training.

### Model construction

We implemented Keras, a high-level API from Tensorflow, to construct and train our deep learning model. The model architecture was based on three CNN blocks. Each CNN block concluded a convolutional layer [filters: (64,32,16), kernel_size: (8,8,8)] with a rectified linear activation function (ReLU), a batch normalization layer, and a dropout layer [rate: (0.5,0.5,0.5)] (Supplementary Figure 3). Finally, a fully connected layer (unit: 1) with a sigmoid activation function produced output values between 0 and 1. We set the batch size of the validation dataset we had produced to 1000. We focused on the validation loss every epoch during model training, and we then stopped training when the training process was stable and the validation loss was no longer decreasing. The model with the lowest validation loss was saved as the best model.

## Results

We used our testing dataset to compare the performance of our model with the following state-of-the-art predictors: AVPpred, AntiVPP1.0, Meta-iAVP, and FIRM-AVP. Table 1 lists the results of this comparison. All the predictors, including that of our model, performed poorly (all accuracy levels at approximately 0.5) after training on the 2012 dataset. Many AVPs have been discovered or synthesized in recent years, so the 2012 dataset is no longer sufficient to represent all AVP features. This caused the classifiers to be trained on incomplete positive data, which resulted in poor performance on the testing dataset. The last two models trained by the datasets we collected performed better, particularly in identifying AVPs (accuracy > 0.8). Our final model (AI4AVP) trained on GAN augmentation data exhibited better performance than the model trained only with real AVP data. GAN augmentation allowed all negative data to be applied to model training, improving the robustness of the classifier for peptide identification.

**Table 1.** Results of our model and other predictors. *2012 datasets were collected by the authors of the AVPpred study with 506 positive and negative data points. **Our dataset without GAN data augmentation (2,641 positive 2,641negative data points). ***Our dataset with GAN data augmentation (16,995 positive and 16,995 negative data points).

| Training dataset | Predictors | Accuracy | Precision | Sensitivity | Specificity | MCC |
| --- | --- | --- | --- | --- | --- | --- |
| 2012 datasets* | AVPpred | 0.55 | 0.61 | 0.29 | 0.82 | 0.12 |
| 2012 datasets* | AntiVPP1.0 | 0.56 | 0.58 | 0.44 | 0.68 | 0.13 |
| 2012 datasets* | Meta-iAVP | 0.55 | 0.56 | 0.49 | 0.61 | -0.04 |
| 2012 datasets* | FIRM-AVP | 0.48 | 0.48 | 0.47 | 0.49 | 0.10 |
| 2012 datasets* | AI4AVP | 0.55 | 0.56 | 0.46 | 0.64 | 0.11 |
| new datasets** | AI4AVP | 0.80 | 0.84 | 0.75 | 0.86 | 0.63 |
| new datasets+<br>GAN dataset*** | **AI4AVP** | **0.84** | **0.84** | **0.85** | **0.86** | **0.68** |

In conclusion, this study developed an AVP predictor, AI4AVP, trained by a more extensive dataset than that used in previous studies. The AI4AVP pipeline is shown in Supplementary Figure 4. Using PC6 encoding and a peptide GAN developed in our previous studies, we achieved data augmentation. This approach allowed all our training data to be utilized and maintained balance between the datasets during model training.

