## Supplements for "Developing an Antiviral Peptides Predictor with Generative Adversarial Network Data Augmentation"

**Supplementary Table 1.** List of the predictors compared with AI4AVP in this study.

| Predictors | Machine learning | Website |
| --- | --- | --- |
| **AVPpred (1)** | Support Vector Machine | http://crdd.osdd.net/servers/avppred/ |
| **AntiVPP1.0 (2)** | Random Forest | https://github.com/bio-coding/AntiVPP |
| **Meta-iAVP (3)** | Ensemble Model | http://codes.bio/meta-iavp/ |
| **FIRM-AVP (4)** | Support Vector Machine | https://msc-viz.emsl.pnnl.gov/AVPR/ |
| **AI4AVP** | **Deep learning** | **https://symbiosis.iis.sinica.edu.tw/AI4AVP/** |

**Supplementary Table 2.** List of databases we collected data from.

| Database | Label | Website |
| --- | --- | --- |
| **APD3 (5)** | Positive | https://wangapd3.com/main.php |
| **DRAMP (6)** | Positive | http://dramp.cpu-bioinfor.org/ |
| **YADAMP (7)** | Positive | https://webs.iiitd.edu.in/raghava/satpdb/catalogs/yadamp/ |
| **DBAASP (8)** | Positive | https://dbaasp.org/home |
| **CAMP (9)** | Positive | http://www.camp3.bicnirrh.res.in/index.php |
| **AVPdb (10)** | Positive | http://crdd.osdd.net/servers/avpdb/ |
| **Swiss-Prot (11)** | Negative | https://www.uniprot.org/ |

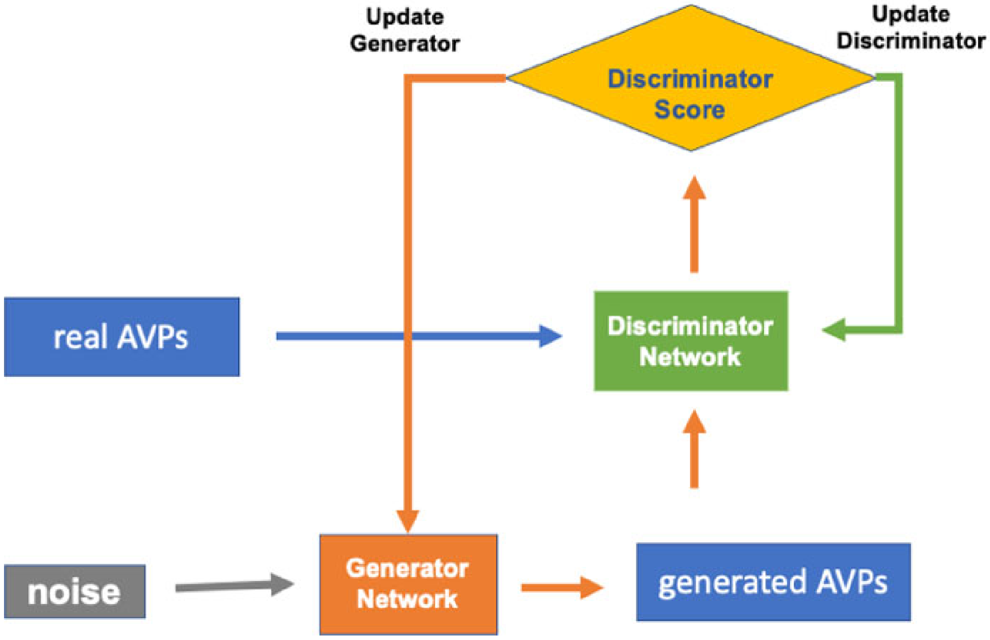


**Supplementary Figure 1.** GAN model for AVP generation. GAN comprises two neural networks: the generator and the discriminator. The generator takes noise as its input and generates AVPs (fake AVPs). The discriminator is trained to distinguish real AVPs from generated AVPs. The purpose of the training process is to make the generator strong enough to produce AVP-like peptides that the discriminator mistakes for real AVPs.

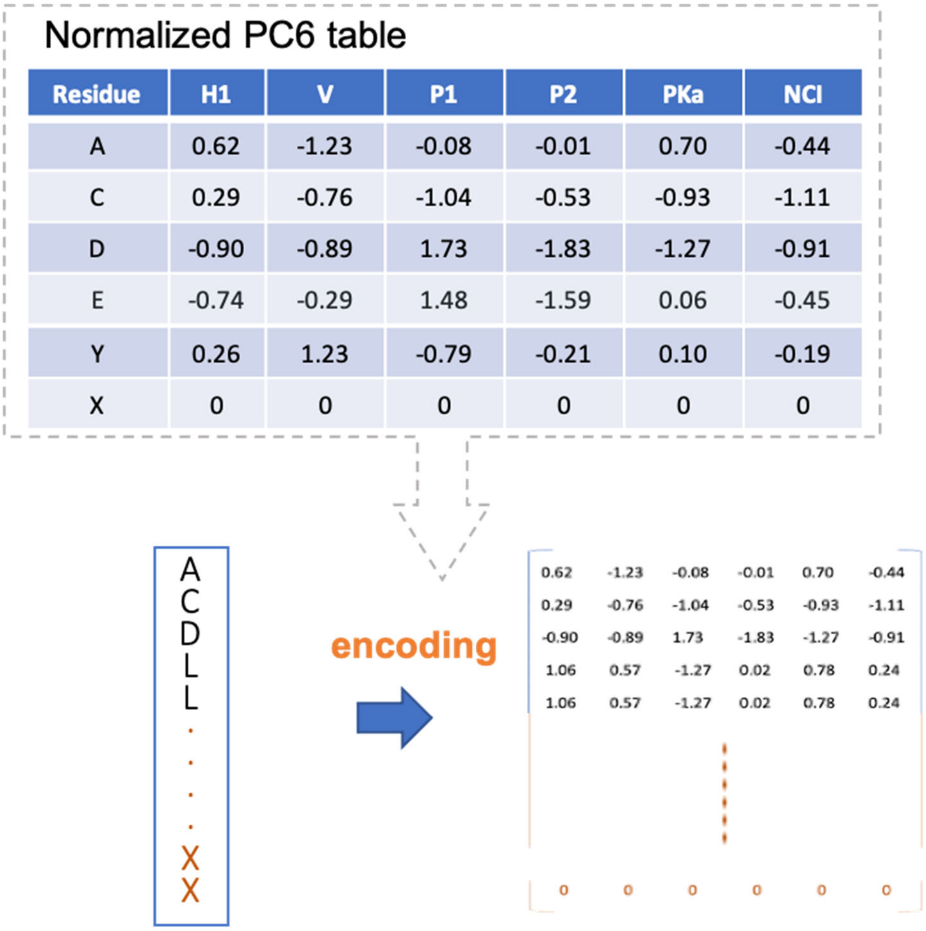

Normalized PC6 table

| Residue | H1 | V | P1 | P2 | PKa | NCI |
|---|---|---|---|---|---|---|
| A | 0.62 | -1.23 | -0.08 | -0.01 | 0.70 | -0.44 |
| C | 0.29 | -0.76 | -1.04 | -0.53 | -0.93 | -1.11 |
| D | -0.90 | -0.89 | 1.73 | -1.83 | -1.27 | -0.91 |
| E | -0.74 | -0.29 | 1.48 | -1.59 | 0.06 | -0.45 |
| Y | 0.26 | 1.23 | -0.79 | -0.21 | 0.10 | -0.19 |
| X | 0 | 0 | 0 | 0 | 0 | 0 |



**Supplementary Figure 2.** PC6 protein-encoding method. As indicated in the figure, the peptide was padded to a length equal to that of the others in the dataset then replacing every amino acid to normalized 6 physiochemicalpropetry values.

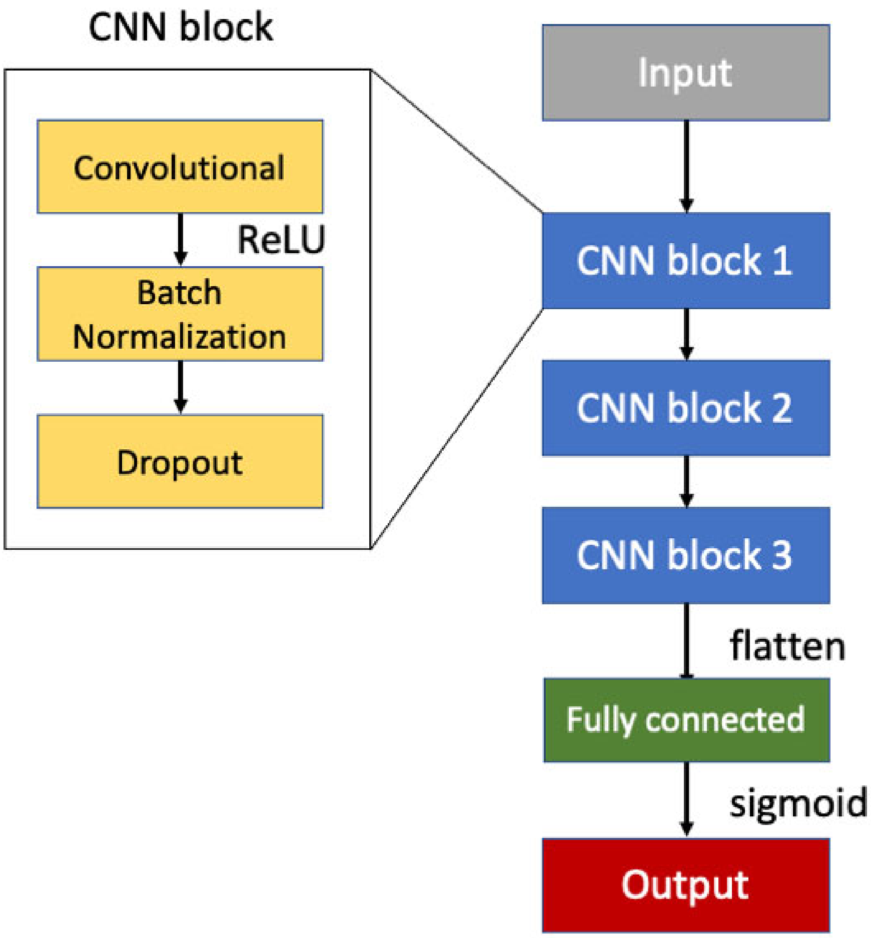


**Supplementary Figure 3.** AI4AVP model architecture visualization. The encoded peptide serves as input to the following three CNN blocks. The fully connected layer with a sigmoid activation function transforms the vector into a value between 0 and 1 as the output (prediction result).

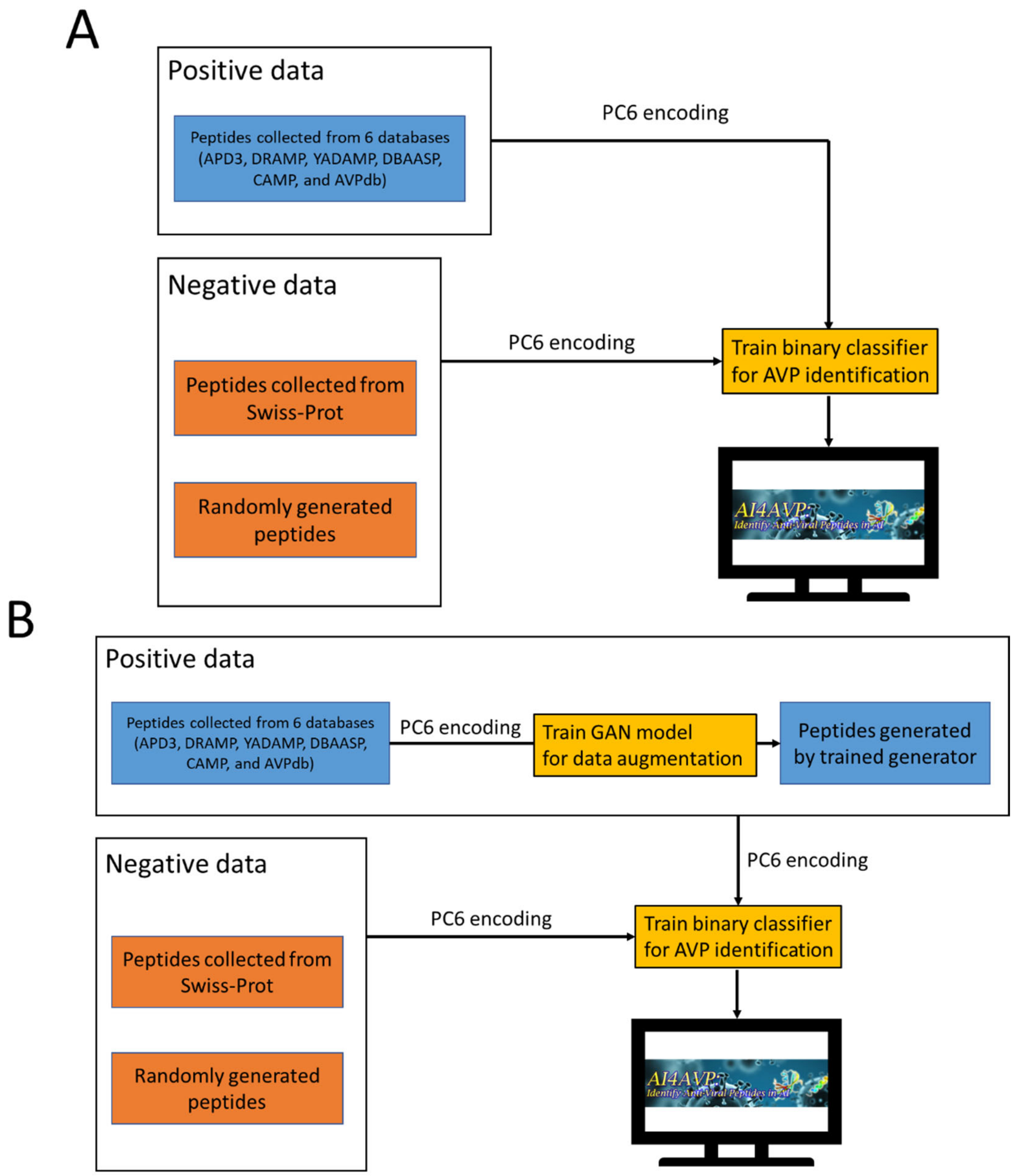


**Supplementary Figure 4.** Pipeline of AI4AVP development. The difference between the models shown in Panels A and B is the dataset. (A) Panel A shows the model trained on a positive dataset consisting of only real AVPs. To balance the positive and negative datasets, the number of non-AVPs was limited. (B) Panel B shows the model trained using real AVPs plus generated AVPs. We applied GAN for data augmentation to utilize all negative data in model training.
